# Occurrence of Lymphedema in Wild-Caught Anurans

**DOI:** 10.1101/336131

**Authors:** Malcolm L. McCallum

## Abstract

Lymphedema is a condition in which the lymph hearts fail to pump fluid from the lymph sacs of anurans and other amphibians. This causes the sacs to fill with fluid and provide the frog with balloon-like swellings or over-all appearance. The condition has previously been connected with various diseases including tadpole edema virus and chytrids. I observed lymphedema in six anuran species (*Acris blanchadi*, Anaxyrus fowleri*, Hyla squirrela*, Pseudacris streckeri illinoensis*, Rana sylvatica, Rana sphenocephala\** [species with * are species records for lymphedema]).

## Introduction

Reports of lymphedema-like disorders in anurans are rare in the literature. This disorder, first described in 1915 (Moore, 1915) has been identified as one of the most common abnormalities in captive bred *Xenopus*. This lymphatic dysfunction most commonly results before or just after metamorphosis. It is characterized by extreme inflammation of tissues giving the frog a ballooned appearance. Apparently the lymph hearts fail to drain the lymph sacs and do not reach maturity (Elkan, 1976). This condition may be an inherited condition due to a recessive gene (Uehlinger, 1965, 1969). Subcutaneous lymph sac structure has been described in some anurans (Carter, 1979).

Gilbert (1942) reported abnormal inflation of the lymph sacs in a pair of male wood frogs (*Rana catesbeiana*) from New York. Their submandibular, ventral, and lateral lymph sacs were filled with air causing them to float helplessly and be incapable of submerging. A similar problem occurs in Illinois Chorus Frogs (*Pseudacris streckeri illinoensis*) in Arkansas (McCallum et al., 2001). This disorder can be confused with the build-up of fluid characteristic of lymphedema.

Captive *Xenopus laevis* are known to develop a lymphedema termed “hydrops” when raised in water with low salinity (Schwabacher and Elkan, 1952). Tadpoles and froglets with hydrops possess inflamed dorsal, lateral, ventral, femoral, and tibial lymph sacs. Wild-caught hydropsic bullfrog (*Rana catesbeiana*) tadpoles have been reported with lymphedema of only the femoral and crural lymph sacs (Hintz, 1963). A similar condition was also reported in overwintering tadpoles and frogs (*Rana mucosa*), (Bradford, 1984).

Iridoviruses known from amphibians are classified as polyhedral cytoplasmic amphibian viruses (PCAV). One of these groups, tadpole edema virus (TEV), mimics subcutaneous hydrops (Marcus, 1981). This virus is transmitted in the water and causes mortality in newly metamorphosed frogs. Its symptoms include subcutaneous edema, petechial hemorrhages, and necrosis in liver, kidney, gastrointestinal tract, and skeletal muscle (Faeh et al., 1998). Wolf et al. (1968) performed histopathologic analysis of edematous *R. catesbeiana* tadpoles identifying a causal agent, the tadpole edema virus (TEV). They later isolated TEV from apparently healthy *R. catesbeiana* though it could not be isolated from either *Anaxyrus* or *Scaphiopus* from the original source pond. TEV was isolated from *R. catesbeiana* tadpoles from Wisconsin, Alabama, West Virginia, North Carolina, and along the Mississippi River in Arkansas (Wolf et al., 1969). However, edema has also been attributed to the bacterium *Flavobacterium indologenes* (Olson et al., 1992) and the fungus *Batrachochytrium dendrobatidis* (Martel et al., 2011).

*Rana catesbeiana* tadpoles appear to be very sensitive to TEV (Wolf et al., 1969), but nothing is known about susceptibility of *A. crepitans* to this pathogen. They found that if the virus was added to water containing 18 day-old *R. catesbeiana* tadpoles, that death ensued beginning after 5 days of exposure. Mortality peaked at 53% by the 7^th^ day of this study. Forty-nine-day-old tadpoles showed first signs of mortality on the 5^th^ day after exposure, with mortality peaking at 37% on the 9^th^ day. All tadpoles were dead by the 13^th^ day. Clear-cut histopathology included necrosis of the liver, kidneys, and digestive tract. Liver necrosis was identified by the second day after exposure. Kidney necrosis and edema was expressed after the 4^th^ day. By the 5^th^ day the digestive tract started to show necrosis.

Because lymphedema is closely associated with TEV it is important to document and report its occurrence in regions where this condition has not previously been reported.

## Materials and Methods

Observations of lymphedema in *Acris crepitans, Pseudacris streckeri illinoensis, Anaxyrus fowleri*, and *Hyla chrysoceles* were noted as reported. Most specimens were released, but others were anesthetized with dilute chloretone, fixed in 10% formalin, preserved in 70% ethanol and deposited in the Arkansas State University Museum of Zoology herpetological collection (ASUMZ), Illinois Natural History Survey collection (INHS), or the Louisiana State University at Shreveport Museum of Life Sciences (LSUSMF).

## Results and Discussion

Incidence of lymphedema among wild-caught anurans was relatively low (Table 1), but was observed in 4 of 5 species in this study. Wood Frogs (*Rana sylvatica*) are a charismatic species of frog common in much of North America. They breed in explosive choruses over a few nights in late winter to early spring. The incidence in Wood Frogs was associated with a die-off of frogs during the breeding chorus in the Sylamore District of the Ozark National Forest in Arkansas (Trauth et al., 2000). In Wood Frogs, lymphedema ranged from the entire torso to localized single lymph sacs on the legs or patches on the back (Fig. 1).

**Figure 1.**
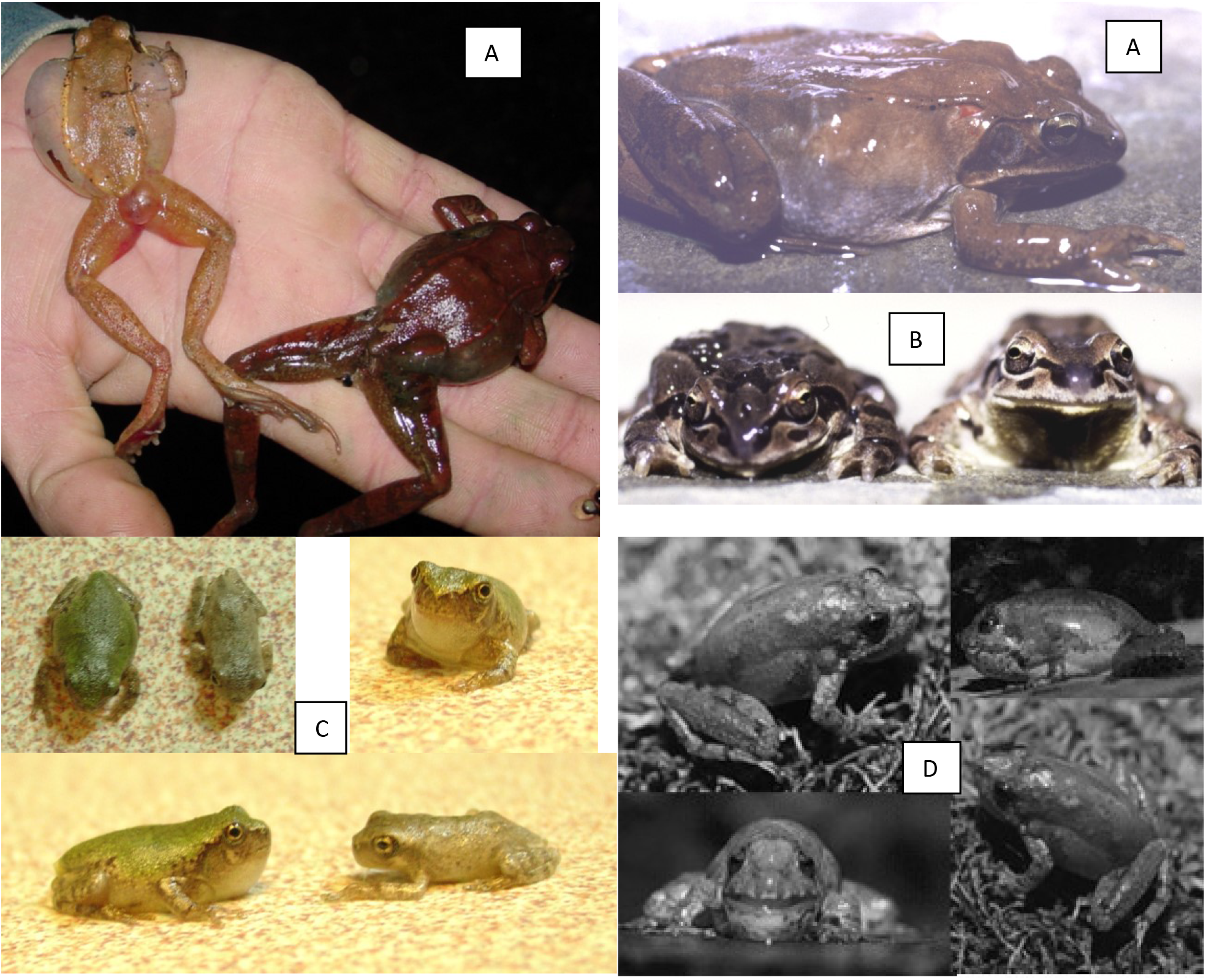
Lymphedema in wild-caught anurans. A) Wood Frog (Rana sylvatica) note: USGS Wildlife Disease Laboratory found no sign of disease (Green Pers. Comm). B. Illinois Chorus Frog (*Pseudacris streckeri illinoensis*). C. Squirrel Tree Frog (*Hyla squirrela*). D. Blanchard’s Cricket Frog (*Acris blanchardi*).

**Table 1.**
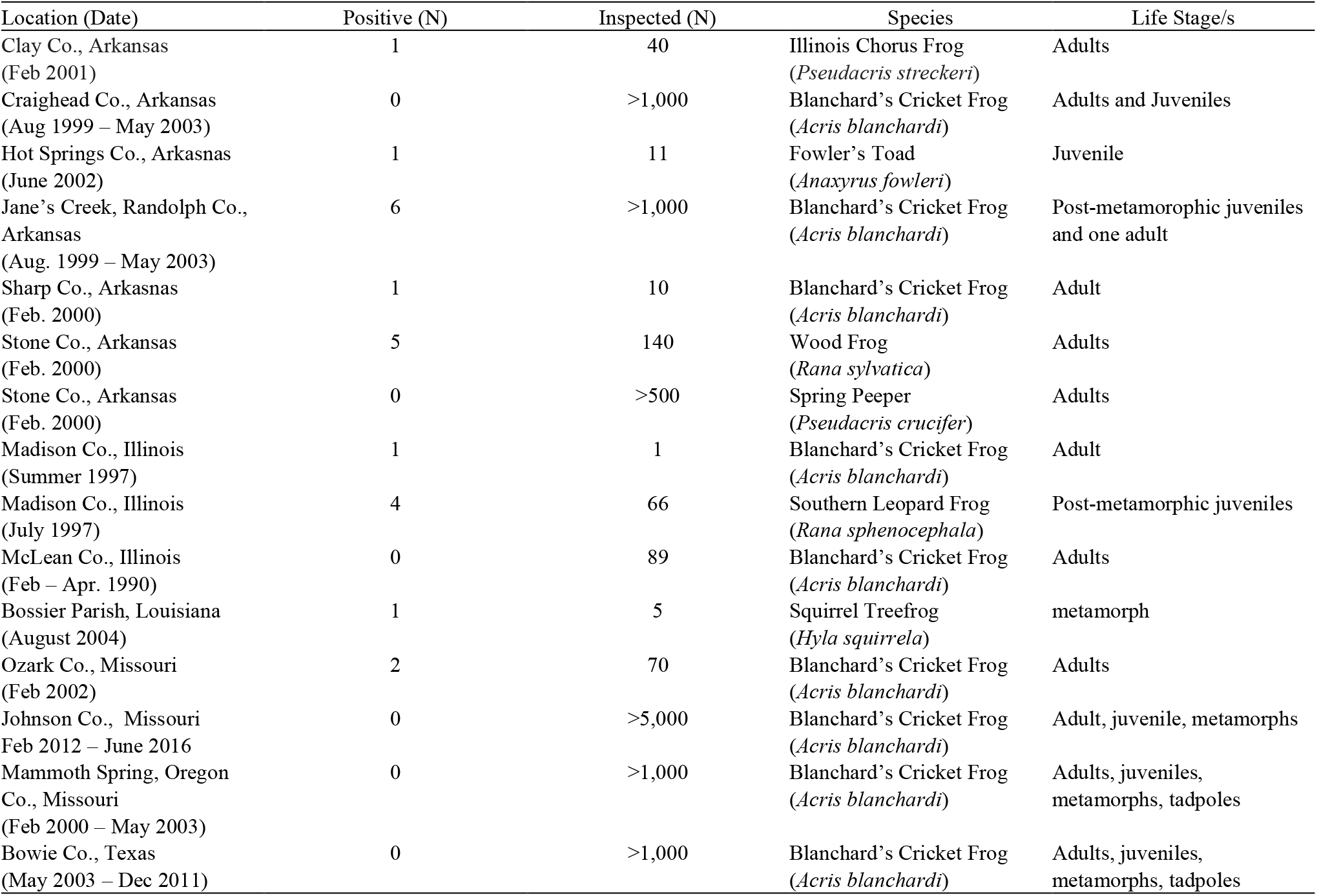
Surveys of amphibians with incidences of lymphedema.

The Illinois Chorus Frog (*Pseudacris streckeri illinoensis*) is a small fossorial frog that occurs in Arkansas. The range of Illinois Chorus Frogs in Arkansas has severely contracted, most likely due to precision leveling of the farmland in which they reside (McCallum and Trauth, 2002; Trauth et al., 2006). I observed a single frog with lymphadema of the right dorsal lymph sac (Fig. 1). This individual appeared normal in movements and activity although it did not appear to be calling or acting as a satellite to other calling males (McCallum et al., 2003). Illinois Chorus Frogs were also observed with frostbite scars (n = 10), red inguinal pustules (n = 2), a dysfunctional vocal sac (n = 1), and a missing forelimb (n = 1), (McCallum et al., 2006; McCallum et al., 2001).

The Southern Leopard Frog (*Rana sphenocephala*) is a generic frog species easily confused with several other species of Leopard Frogs whose ranges they overlap. They are common to creeks, grasslands, forests, and ponds throughout the Southeastern United States. Among 66 post-metamorphic Southern Leopard Frogs that were examined, a few (est. 3-4) were observed with lymphedema. However, the exact numbers were not recorded among malformed frogs in that study (McCallum, 1997), nor were possible incidences of partial lymphedema noticed because I was not aware of the condition or its connection with diseases that early in my career.

Fowler’s Toad (*Anaxyrus fowleri*) is an average sized toad occurring across the eastern half of the United States. A single metamorph had lymphedema. It seemed otherwise normal, except that it had difficulty moving with the disorder. No Spring Peepers (*Pseudacris crucifer*) or Gray Treefrogs (*Hyla versicolor* or *H. chryscocelis*) were observed with the condition despite occurring in locations where other species had it (Wood Frogs, Southern Leopard Frogs, Blanchard’s Cricket Frogs).

The Squirrel Treefrog (*Hyla squirella*) is a common arboreal frog of the Southeastern United States, although it was unknown to the Shreveport, Louisiana area at the time of this observation. We collected five near-metamphosed tadpoles at Loggy Bayou Wildlife Management (Bowie Co., Louisiana, USA) and placed them in a 20 L aquarium containing water from the collection site. They appeared morphologically normal. Upon metamorphosis, one developed lymphedema, the rest were normal. This frog died seven days later.

Blanchard’s Cricket Frog (*Acris blanchardi*) is a small non-arboreal hylid of the eastern half of North America that can reach extraordinarily high populations on stream banks in the Ozarks and elsewhere (McCallum and Trauth, 2004; McCallum et al., 2011) which facilitates some of its unique anti-predator behaviors (McCallum, 1999; McCallum, 2011). Its taxonomic status with *Acris crepitans* is currently subject to debate (McCallum, 2003; McCallum and Trauth, 2006). Anatomical abnormalities in Blanchard’s Cricket Frog from Arkansas have increased over the past 43 years (McCallum and Trauth, 2003) and its reproductive and growth strategies (McCallum and Trauth, 2007; McCallum et al., 2011) appears to place it at threat from climate change (McCallum, 2010). I observed large numbers of living Blanchard’s Cricket Frog in five different states (Arkansas, Illinois, Louisiana, Missouri, and Texas). Addtionally, I inspected hundreds of preserved frogs from South Dakota, Nebraska, Georgia and Florida during other studies (McCallum, 2003; McCallum et al., 2011) although it was difficult to assess lymphedema in preserved specimens that had not also been seen alive. I observed more cases of lymphedema in Blanchard’s Cricket Frog (Fig. 1) than in other species, but this was probably more due to sampling effort. Lymphedema was relatively rare (Table 1). These frogs were less mobile compared to non-afflicted individuals which would place them at a significant threat to predation, including from adults (McCallum et al., 2001), because their anti-predator behavior depends on rapid movement (McCallum, 2011).

Except in cases with minor lymphadema consisting of single or partial lymph sac inflammation, the disorder can be very debilitating. Frogs with extreme swelling have difficulty moving, jumping, and often remain motionless much of the time. If the disorder is caused by TEV or chytrids then it would constitute a possible method of dissemination if metamophs emerged unable to escape hungry adult frogs of the same and other species. Although the condition of lymphedema has been known for many years, it is surprising that more studies have not been conducted in the wild considering its association with amphibian pathogens and the rapidly accelerating extinction of amphibians we have already observed (McCallum, 2007; McCallum 2015).

